# Combating Chagas Disease Through Inhibition of Tiam1, a Rho GTPase Guanine Nucleotide Exchange Factor

**DOI:** 10.1101/040121

**Authors:** Chirag Krishna, Li Xie, Celine DerMarderossian, Philip E. Bourne

**Affiliations:** Department of Biology, University of California San Diego, La Jolla, California, United States of America; Skaggs School of Pharmacy and Pharmaceutical Sciences, University of California San Diego, La Jolla, California, United States of America; The Scripps Research Institute, Department of Immunology and Microbial Science, La Jolla, California, United States of America

## Abstract

Chagas disease is a major cardiovascular affliction primarily endemic to Latin American countries, affecting some ten to twelve million people worldwide. The currently available drugs, Benznidazole and Nifurtimox, are ineffecteive in the disease’s chronic stages and induce severe side effects. In an attempt to improve this situation we use an *in silico*drug repurposing strategy to correlate drug-protein interactions with positive clinical outcomes. The strategy involves a protein functional site similarity search, along with computational docking studies and, given the findings, a phosphatidylinositol (PIP) strip test to determine the activity of Posaconazole, a recently developed antifungal triazole, in conjunction with Tiam1, a Rho GTPase Guanine Nucleotide Exchange Factor. The results from both computational and *in vitro*studies indicate possible inhibition of phosphoinositides via Posaconazole, preventing Rho GTPase-induced proliferation of *T. cruzi*, the etiological agent of Chagas Disease.

**Synopsis:** While Benznidazole and Nifurtimox are effective in the acute phase of Chagas’ disease, their inefficacy in the disease’s chronic phase and severe side effects motivate the search for more novel treatment pathways. Antifungal triazoles have shown promise in murine models of Chagas. Using a computational platform, the authors find that these drugs may be used to combat Chagas Disease through a previously unknown mechanism. If correct, this model could be used to motivate the production of new vaccines for Chagas. Furthermore, the model demonstrates the efficacy of computational testing in the preliminary search for effective drug candidates.

## Introduction

Chagas Disease is a major cardiovascular affliction primarily endemic to Latin American countries, affecting millions of inhabitants of Mexico, Central America, and South America. The proliferation of its etiological agent, *Trypanosoma Cruzi*, causes fatal cardiac and intestinal disorders in its chronic phase. Available drugs such as Benznidazole and Nifurtimox are effective in treating acute Chagas, yet their limited efficacy in the chronic phase and toxicity are motivation to investigate safer, more effective compounds [1]. Posaconazole, a recently developed antifungal triazole, is a new experimental compound that is known to be effective in treating mice infected with chronic Chagas. Posaconazole is highly potent and has better pharmacological properties than current drugs Benznidazole and Nifurtimox. Furthermore, its status as the only drug in its class to be tested in clinical trials makes it a more attractive candidate for further research. By identifying relevant off-targets for posaconazole within the human proteome, it may be possible to repurpose the drug for increased effect against *Trypanosoma cruzi* and less toxicity for humans. Subsequent chemical modification could increase its affinity for the intended target, and decrease its affinity for other off-targets that may cause adverse side effects. Furthermore, studying the binding of approved compounds to off-targets may also result in the creation of a drug outside the scope of its patent, hence increasing the likelihood of its development as a therapeutic product.

Previously we detailed an algorithm for comparing a template ligand binding site to other putative binding sites across fold and functional space as exemplified by the proteome of an organism [3]. The original binding site of Posaconazole is found to target 14-alpha sterol demethylase (PDB ID: 2X2N), an enzyme responsible for the conversion of lanosterol to ergosterol, a key component of fungal cell membranes. To identify similar binding sites for Posaconazole, we scanned its original binding site against the human proteome. Our algorithm predicts with high probability that Tiam1, a guanine nucleotide exchange factor of Rho GTPases, is a potential binder of Posaconazole. Tiam1 facilitates activation of the Rac1 GTPase [4], which, through an undefined mechanism, is correlated with *T. cruzi* amastigote cell invasion [5]. The entry of *T.cruzi* trypomastigotes into the cell is known to strongly stimulate the formation of lipid products; in particular, phosphatidylinositol-3-phosphate (PI3P) [6]. The loss of PI3P binding to the pleckstrin homology (PH) domain of Tiam1 results in decreased Rac1 expression [4], suggesting that inhibitors of this binding site could slow proliferation of *T. cruzi* amastigotes and thus be of therapeutic value. Therefore, we decided to further examine the interactions between Posaconazole and Tiam1. Using protein-ligand docking and molecular visualization, we compared the Posaconazole-Tiam1 complex to that of PI3P-Tiam1. Our prediction shows possible competition between PI3P and Posaconazole for Tiam1 binding, which could be exploited for further Chagas disease drug development efforts. Furthermore, we evaluated additional potential lead compounds identified via virtual screening that may display inhibitory activity *in vitro*.

## Methods

### Structural models of the human proteome

Sequences of all PDB [18] structures are mapped to Ensembl [19] human protein sequences (43,738 proteins) using BLAST [20]. A total of 10,730 PDB structures map to 3,158 Ensembl human proteins with a sequence identity above 95%. These 10,730 structures are considered as structural models for the human proteome. They form 2,586 sequence clusters when using a sequence identity of 30%.

### Structural coverage of the druggable human proteome

The existing druggable human proteome is determined by mapping Ensembl [19] human protein sequences against all sequences of drug targets from Drugbank [21] using BLAST [20]. Homologous sequences from the human proteome, with e-values less than 0.001, constitute the druggable human proteome—a total of 13,865 human proteins corresponding to 2,002 unique drug targets. Among the 10,730 human protein structural models, 1,585 belong to the existing druggable human proteome and correspond to 929 unique drug targets. 825 sequence clusters are formed after clustering the druggable structures with a sequence identity of 30%. One structure is randomly selected from each of the clusters to constitute a representative set of druggable structures. These structures represent approximately 10% of the complete druggable human proteome and 40% of existing drug targets.

### Ligand binding site similarity

Protein structures are represented by Delaunay tessellation of Cα atoms and characterized with geometric potentials [22]. The similar residue clusters for any protein to a ligand binding site are detected with a SOIPPA algorithm [30]. To evaluate the *p*-value for the similarity score calculated from the site comparison method, we estimate the background distribution using a non-parametric method. First, the drug target of interest is compared against the 825 representative sets of human druggable structures. We remove those hits that are in the same fold as the query because they will probably be true positives. Then a kernel density estimator is used to estimate the background probability distribution of the binding site alignment scores. A Gaussian kernel with fixed bandwidth is used. The optimal bandwidth is estimated from the data by using a least square cross-validation approach [23]. Finally, this estimated density function is used to calculate a *p*-value for the particular pair of ligand sites being compared.

### Protein-Ligand Docking

Protein-ligand docking is conducted using Autodock Vina. The default parameter settings were applied, with the exhaustiveness set at 8 during docking, corresponding to the most extensive conformational search over the protein structure. The structures used in this study were downloaded from the RCSB PDB [18]. The PDB id of the original Posaconazole site (14-alpha sterol demethylase) was 2X2N. The id of Tiam1 was 3A8N. The PDB ligand id for Posaconazole was X2N.

### PIP Strip Testing

The PIP membrane is wet in PBS/Tween for several minutes and incubated with blocking buffer, 0.1% PBS/Tween in addition to 3% nonfat dry milk for one hour at room temperature. The blocking buffer is discarded, drug and protein are added at 0.25-1 ug/ml at room temperature. The membrane is washed using 0.1% PBS/Tween for 3 to 5 minutes. The membrane is incubated with primary antibody in blocking buffer for 1 hour at room temperature. The membrane is washed with 0.1% PBS/Tween for 3 to 5 minutes. The membrane is incubated with secondary antibody in blocking buffer for 1 hour at room temperature, and washed again with 0.1% PBS/Tween. The protein is then detected.

## Results

First, 10,730 structures were selected from the RCSB Protein Data Bank [29] by mapping sequences of PDB structures to Ensembl human proteins [24] using a sequence identity of over 95%. Of the 10,730 structures, 929 are unique drug targets (see Methods). Using a sequence identity cutoff of 30%, 825 structures are chosen - a set representing 10% of targetable human proteins. By taking homology models into consideration, we estimate that the structural coverage of the human genome is approximately 40%. While the set is far from complete, several proteins of interest were identified as relevant off-targets in the context of Chagas disease.

This set of 825 representative structures was scanned for structural similarity against the ligand binding site of 14-alpha sterol demethylase (PDB id: 2X2N) [7]. Our previous studies have indicated that structures with similarity defined by a *p*-value < 0.001 are most significant. Our sequence order independent profile-profile alignment (SOIPPA) algorithm yielded 80 structures within the *p*-value threshold. Of these, 25 were, or were derived from, Cytochrome P450 proteins. Cytochrome P450s are known to be promiscuous drug binders and are already very well characterized in the context of drug metabolism [8]. Because this study was focused on identifying novel uses for Posaconazole, these results were discarded. For the remaining 55 proteins, a literature review was conducted to determine the most likely relevance to Chagas disease. Additionally, we utilized the Kyoto Encyclopedia of Genes and Genomes (KEGG) [25, 26] to examine the Chagas related protein network, and to relate nested networks within that of Chagas (i.e., phosphatidylinositol signaling pathway) to proteins from the algorithmic output.

Following this review, it became clear that Tiam1, a Rho GTPase guanine nucleotide exchange factor, was the best candidate for further investigation. Tiam1 is comprised of two domains - the DBL Homology (DH) domain, and Pleckstrin Homology (PH) domain [9]. While the DH domain of Tiam1 is primarily involved in guanine nucleotide exchange (GEF) activity, the PH domain binds several phosphoinositides which modulate the activation of the Rac1 GTPase. Lemmon *et al*. showed that loss of phosphatidylinositol-3-phosphate (PI3P) binding to the PH domain prevented activation of the Rac1 GTPase *in vivo* [4]. It has been suggested that Rac1 activation is correlated with *T. cruzi* amastigote invasion [5]. Because the entry of *T. cruzi* trypomastigotes into the cell is known to strongly stimulate the formation of PI3P, an inhibitor of the PI3P binding site in the PH domain of Tiam1 could slow or prevent the proliferation of *T. Cruzi* amastigotes. As Tiam1 was predicted to retain a high degree of structural similarity to the original binding site of Posaconazole (*p*-value 1.47 x 10^−7^), we decided to computationally study the binding site of Posaconazole in Tiam1, and compare it to that of PI3P in Tiam1.

For this study, Autodock Vina was selected to perform molecular docking based of its improved accuracy and performance over other docking programs, such as Autodock 4. Vina’s functionality is grounded in machine learning methods, rather than in an exclusive focus on traditional physics concepts (i.e., Lennard-Jones Potential and Coulomb Energy) [10]. Vina performs rigid docking of a ligand to a given protein, and outputs a variety of different conformations accompanied by RMSD values and free energy scores (kcal/mole). These various conformations can be compared to the original binding pose to determine the likelihood of a particular pose *in vitro*.

We first docked PI3P to the PH domain of Tiam1 to determine its binding site. Theoretically, the binding site of PI3P should correspond to the site of interest when evaluating potential inhibitors. The search space was set over the entire Tiam1 structure (DH and PH domains) to determine alternative binding sites for PI3P. However, the docked conformations agree with results articulated by the Lemmon *et al* group. PI3P appears to bind preferentially to the PH domain of Tiam1 - of the nine prioritized poses output by Vina, poses 1-5 and 7 bound at the same location in the PH domain. While low docking scores initially suggested the low probability of binding, Lemmon *et al*. noted that the PH domain bound PI3P and other phosphoinositides weakly. Thus, the consensus binding was most important in determining a potential inhibition site for further evaluation.

We next docked Posaconazole to Tiam1, setting the search space over the entire structure. An interesting characteristic of Posaconazole is its varied conformations within 14-alpha sterol demethylase - it is known to adopt both bent and straight conformations. The primary difference between the two conformations is the presence of a single hydrogen bond between the terminal oxygen atom of Posaconazole and the contacting receptor residue in the bent conformation. Of the nine poses output by Vina when bound to Tiam1, seven were in the straight conformation. The Accelrys Discovery Studio 3.1 [27] client was used to monitor the interactions between ligand and receptor. Not surprisingly, the software detected no hydrogen bonding between Posaconazole and Tiam1 in its straight conformations. Pymol [12] was used to monitor specific interactions between Posaconazole and Tiam1. Ligand-receptor contact with distances between 2.0 and 4.0 Angstroms were considered. In each of its straight conformations, Posaconazole displays many hydrophobic and van der Waals interactions, similar to its activity within 14-alpha sterol demethylase. Furthermore, six of the nine conformations output by Vina bound in the same location as PI3P. While the docking scores for Posaconazole are not particularly strong, it seems possible that the molecule may compete with PI3P for binding within the PH domain. This hypothesis is further verified by the fact that Posaconazole is a very large molecule (molecular weight = 700.8 g/mol) and completely blocks PI3P from its binding site. Moreover, Posaconazole and PI3P are chemically different molecules - thus, it is highly unlikely that Posaconazole will induce Rac1 activation *in vivo*. Rather, the competition between Posaconazole and PI3P for binding within the PH domain indicates a role for the former as a Tiam1 inhibitor.

While Tiam1 was the primary receptor of interest, the SOIPPA algorithm yielded several other proteins that may be viable candidates for further Chagas research. To classify these proteins into representative groups, we used the DAVID annotation database (Version 6.7) to cluster the remaining hits into gene-enriched groups [13]. As detailed previously, only proteins with *p*-value < 0.001 were considered. CYP450s were also discarded from the annotation. Accession numbers for these proteins were provided by Uniprot [14]. 23 annotation clusters were generated, each comprised of non-unique elements. To narrow the results further, proteins from sub-clusters with *p*-value < 0.001 were noted. Of particular interest were the human MCAD:ETF E165betaA complex (PDB id: 2A1T) and Human Thioredoxin Reductase 2 (PDB id: 2J3N), due to their involvement in flavin adenine nucleotide activity (FAD), electron transfer, or oxidation-reduction reactions. As an antifungal triazole, Posaconazole disrupts the close packing of phospholipid chains, impairing the function of enzymes involved in the electron transfer system [15]. As prior research points to these proteins as implicated in Chagas disease [16], they certainly warrant further investigation as potential drug targets by Posaconazole or other novel inhibitors.

### *In Vitro* Studies

In order to obtain a preliminary account of Posaconazole’s interaction with Tiam1 *in vitro*, we incubated Posaconazole along with Tiam1 on a standard phosphatidylinositol (PIP) strip containing all eight phosphoinositides. The resulting strip displayed all eight phosphoinositides, including PI3P, blotted out completely, suggesting possible competition between Posaconazole and these molecules within the Pleckstrin Homology domain.

These data from the PIP Strip are not conclusive. Given that Posaconazole is a large molecule, we did not anticipate inhibition of phosphoinositides other than PI3P. This suggests possible binding delocalization of Posaconazole over the PH domain, which may detract from its ability to inhibit PI3P specifically. In future work, more extensive tests should be conducted to better determine Posaconazole-Tiam1 interactions *in vitro*.

## Discussion

Currently available Chagas disease treatments, such as Benznidazole and Nifurtimox, are far too toxic to be considered long term solutions. Their adverse side effects and limited efficacy in the chronic phase are motivation to investigate improved compounds [1]. Posaconazole’s favorable pharmacological properties and documented *in vivo* activity make it an attractive candidate for further review. In this study, we show how our algorithm can produce proteins that are not only structurally similar to a template ligand binding site, but are also relevant to the system of interest. While Tiam1 was the primary focus of this study, the SOIPPA algorithm also predicted that proteins such as the human Thioredoxin Reductase (*p*-value 1.50 x 10^−4^) and the human human MCAD:ETF E165betaA (*p*-value 5.47 x 10^−4^) were structurally relevant. As other work has implicated these proteins in Chagas disease [28], the ability of our model to produce relevant proteins is indirectly validated.

While the docking output presents evidence for possible competitive activity between Posaconazole and PI3P, several shortcomings of modern-day molecular docking must be taken into account. It is well known that proteins are not rigid; rather, they are highly dynamic structures. However, like other docking programs, Autodock Vina does not account for receptor flexibility [10], instead docking small molecules to a rigid structure that is only a “snapshot” of the protein at a given point in time. Furthermore, Posaconazole is a large, hydrophobic molecule that well exceeds Lipinski’s rule of five - its structural characteristics may account for the weaker docking scores attributed to its poses output by Vina. Further studies may utilize molecular dynamics or the relaxed complex scheme to further characterize binding affinities between small molecules and Tiam1, as the aforementioned techniques can provide an ensemble of rigid receptor structures to which molecules can be docked. As the focus of this study was an evaluation of the predictive power of the SOIPPA algorithm and an investigation of possible inhibitors, a single structure was sufficient to determine the possibility of novel competitive behavior.

Our method may serve as an improved first step in the modern systems pharmacology-based drug discovery pipeline. As existing compounds such as Posaconazole are discovered to be effective against various off-targets, medicinal chemists may choose to structurally repurpose these molecules to make them more effective *in vivo*. While our previous studies have focused on reducing unwanted side effects by minimizing off-target binding [17], researchers may wish to optimize Posaconazole to increase its binding affinity for Tiam1 and thus better inhibit PI3P binding.

## Conclusion

Tiam1 is a Rho GTPase guanine nucleotide exchange factor containing a PH domain that binds PI3P. In doing so, it catalyzes the activation of the Rac1 GTPase, which is known to facilitate the proliferation of *T. cruzi* amastigotes, the etiological agents of Chagas disease. Here, we present evidence for a novel use for Posaconazole, a recently developed antifungal triazole that may inhibit PI3P-PH binding and thus slow or prevent the spread of the disease. The need for improved treatments for Chagas and other neglected tropical diseases is dire - a lack of sustained research in this area frustrates attempts to find long-term solutions for these conditions. Our study presents a hypothesis that may guide further Chagas drug discovery projects, specifically in the context of Tiam1 and the Rac1 GTPase. Further studies may consider these proteins, their mechanisms of actions, and related networks more carefully in an effort to expedite the development of a permanent vaccine for Chagas disease.

